# Mechanics of CaMKII-actin networks

**DOI:** 10.1101/308353

**Authors:** Shahid Khan, Kenneth H. Downing, Justin E. Molloy

**Affiliations:** Molecular Biology Consortium, Lawrence Berkeley National Laboratory, Berkeley, CA 94720. USA; Molecular Biophysics & Integrated Bio-Imaging, Lawrence Berkeley National Laboratory, Berkeley, CA 94720. USA; The Francis Crick Institute, London, NW1 1AT, UK

## Abstract

Calcium calmodulin dependent kinase (CaMKII) has an important role in dendritic spine remodelling upon synaptic stimulation. Using fluorescence video microscopy and image analysis, we investigated the architectural dynamics of rhodamine-phalloidin stabilized F-actin networks cross-linked by CaMKII. We used automated image analysis to identify F-actin bundles and cross-over junctions and developed a dimensionless metric to characterize network architecture. Similar networks were formed by three different CaMKII species with ten-fold length difference in the linker region between the kinase domain and holoenzyme hub; implying linker length is not a primary determinant of F-actin binding. Electron micrographs showed that, at physiological molar ratios, single CaMKII holoenzymes cross-linked multiple F-actin filaments in random networks, whereas at higher CaMKII / F-actin ratios filaments bundled. Light microscopy established that random networks resisted macromolecular crowding, with polyethylene glycol mimicking cytoplasmic osmolarity, and blocked ATP-powered compaction by myosin-2 mini-filaments. Importantly, the networks disassembled following addition of calcium calmodulin and were then rapidly spaced into compacted foci by myosin motors or, more slowly, aggregated by crowding. Single molecule TIRF microscopy showed CaMKII dissociation from surface-immobilized G-actin exhibited a mono-exponential dwell-time distribution, whereas CaMKII bound to F-actin networks had a long-lived fraction, trapped at cross-over junctions. Release of CaMKII from F-actin, triggered by calcium calmodulin did not require ATP (hence phosphorylation) and was too rapid to measure with video-rate imaging. The residual bound-fraction was reduced substantially upon addition of an NMDA receptor peptide analogue. These results provide mechanistic insights to CaMKII-actin interactions at the collective network and single molecule level. Our findings argue that CaMKII-actin networks in dendritic spines are stable enough to protect the basal network architecture against physical stress but once CaMKII is disengaged by calcium calmodulin and sequestered by receptors at the synapse; F-actin compaction by myosin motors stabilizes the expanded spine compatible with the recorded times.

## Introduction

The calcium calmodulin dependent kinase II (CaMKII) is a multi-functional, dodecameric kinase assembly. It is ubiquitous in the animal kingdom and has key roles in learning and cardiovascular function. The role in learning has been studied in vertebrate rat models of memory (1, 2) while its effects on dendritic morphology and synaptic localization in the invertebrate nematode, *Caenorhabditis elegans*, have been described (3). CaMKII is important both in long-term potentiation (LTP) and long-term depression (LTD) with the biochemical mechanisms best understood for LTP in hippocampal neurons (4).

The dendritic spine cytoskeleton is composed overwhelmingly of actin. Actin binding by CaMKII and its abrogation by kinase activation is central to spine remodelling during learning and development (5). Our interest is the early phase of LTP that lasts for about three minutes resulting in increased volume of the stimulated spine; characterized as a long-lived “synaptic tag” (6). In addition to its kinase activity, there is accumulating evidence for a direct structural role of CaMKII in dendritic spines (7-9) consistent with its high (> 0.1 mM) abundance (10) and actin binding affinity. Photo-release of caged glutamate has elucidated early LTP dynamics by triggering calcium transients (11) to activate multiple kinases via CaMKII within seconds (12). Changes in F-actin content measured post-stimulation by photo-activation of PaGFP-actin revealed dynamic (< 1 minute) and stable (~ 17 minute) pools with CaMKII required for formation of the stable pool (13). Single particle tracking (SPT) of CaMKII reports three kinetic sub-populations in spines; of which the major immobile subpopulation interacts with the actin cytoskeleton (7, 14). CaMKII associates with actin stress fibres in vivo (8, 15) and F-actin bundles in vitro (9, 16, 17). The binding of rat CaMKII β isoform (β_Rat_) to globular actin (G-actin) (2.4 µM affinity) (17) and neuronal F-actin cytoskeleton (18) is abolished by calcium bound calmodulin (“calcium-calmodulin”); noted to be within five seconds in the latter study.

Myosin IIb activity is also essential for early LTP (19-21). Myosin IIb at the spine neck (22) may regulate traffic between the spine head and dendritic shaft (23). Myosin IIb phosphorylation regulates the size of the post-synaptic receptor patch known as the post-synaptic density (PSD) (24, 25) while knockdown of a sarcomeric myosin II isoform, MyH7B, enlarges spine heads (26). Upon synaptic stimulation, the spine head volume rapidly increases 3-fold, then compacts over ~200 seconds to a stable end-state (~1.5x the pre-stimulated basal volume) (12) (Figure 1). An attractive idea is that the compaction phase is driven by MyH7B, tethered to PSD-associated SynGap / PSD-95 / NMDA receptor complexes (27), or another PSD-localized Myosin IIb. Myosin motor activation could involve mini-filament formation following myosin light-chain kinase (MLCK) activity triggered by the calcium pulse (28) upon synaptic stimulation.

**Figure 1:**
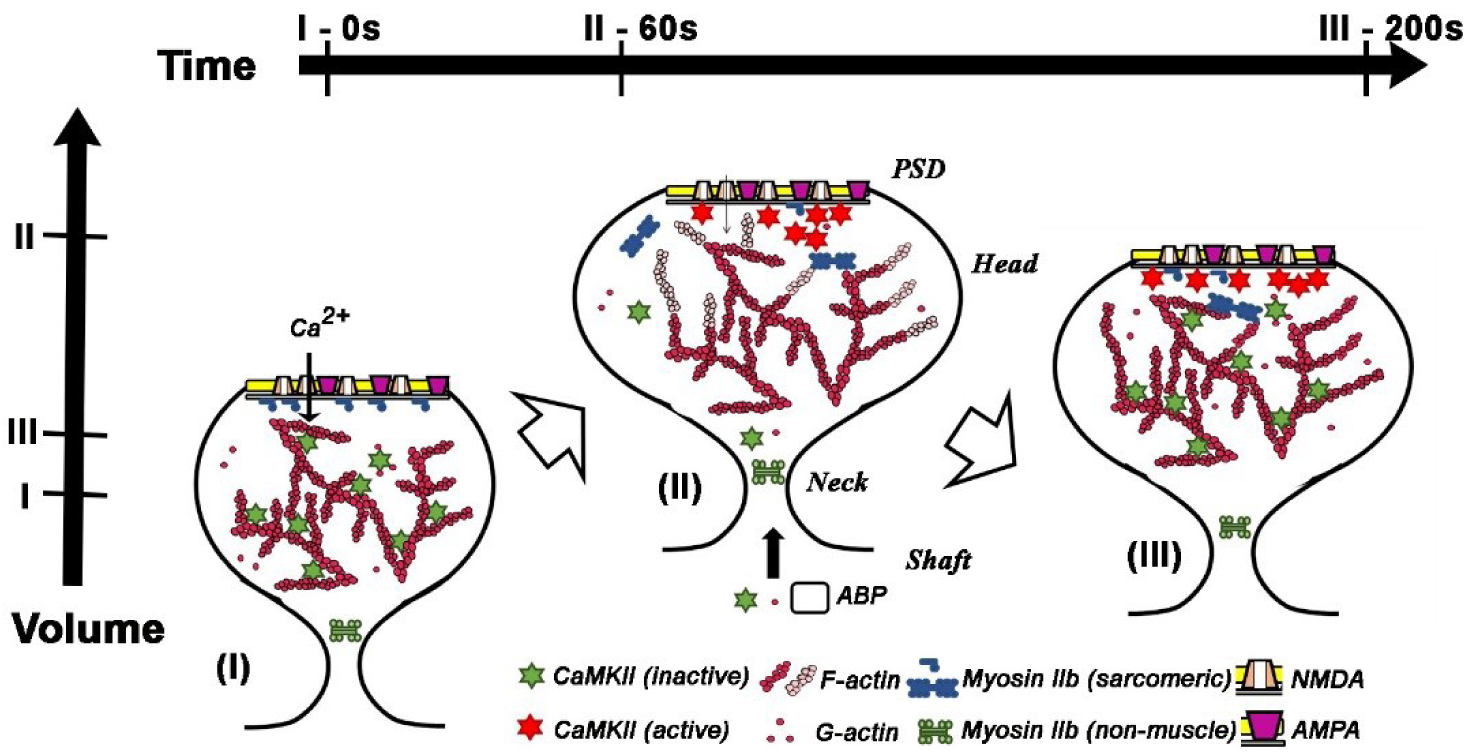
Stimulus-dependent remodelling of the spine cytoskeleton. Spine morphology is shown in its initial **(i)** maximally expanded **(ii)** and stable end states **(iii)** following synaptic stimulation. **(i)** –> **(ii)**: Transient (sub-second) NMDA receptor calcium influx triggers calmodulin mediated CaMKII activation, dissociation from actin and sequestration to the PSD. Actin polymerization, stimulated by increased G-actin levels together with diffusion from the shaft of actin-binding proteins with severing activity (e.g. cofilin), expands the actin cytoskeleton. The activated CaMKII initiates phosphorylation cascades to biochemically orchestrate this expansion. PSD-localized myosin IIb is activated by the calcium pulse, possibly by mini-filament formation mediated by MLCK kinase. **(ii)** –> **(iii)**: As intracellular calcium returns to basal levels compaction of the expanded cytoskeleton by myosin is completed and it is stabilized by attachment of CaMKII that has entered from the shaft. The phospho-relays triggered by the persistently activated PSD CaMKII recruit receptors and structural components as the sustained volume increase leads to eventual increase in PSD size. Horizontal bars denote time stamps for the states while the vertical bars mark relative spine head volumes.

A quantitative model (29) based on transcriptome analysis (30) and live cell imaging of kinases (RhoA, Cdc42) and actin binding proteins (ABP – e.g. cofilin / Arp2/3) mobilized by CaMKII (6, 31) accounts for the observed spine volume changes. Model simulations of actin polymerization triggered by the CaMKII mediated phosphorylation cascades followed by compaction by myosin show spine volume changes are robust to changes in individual components of the biochemical circuity as assessed by sensitivity analysis (32). Current understanding of the spatiotemporal relationships between CaMKII, actin, myosin and spine volume is schematized in **Figure 1**.

In this study, we investigated the architecture of CaMKII cross-linked F-actin (CaMKII-actin) networks (both gels (3D) and surface-attached F-actin layers (2D)) and their response to mechanical stress caused by macromolecular crowding and myosin motor forces. Our in vitro measurements provide information about maintenance of the initial and end states of the stimulated spine as well as the transition between them. Responses of actin networks to crowding agents and myosin motors have been characterized previously (33-37). Here, we study the modulatory role of CaMKII and calcium-calmodulin under conditions that are physiologically relevant to dendritic spines.

Actin-CaMKII interactions are reported to vary with CaMKII isoform (38). Although the canonical CaMKII kinase (KD) and association domains (AD) are conserved, the intervening linker region (β_Rat_316-405) varies in length and sequence between isoforms (39). The conserved AD forms the holoenzyme structural hub. Auto-inhibition of kinase activity is a common strategy; with the ATP binding site altered and blocked by a pseudo-substrate regulatory sequence (β_Rat_275-segmented into R_1_275-291,R_2_292-298,R_3_308-315) (40). Calcium-calmodulin binds to an R_2_ IQ-10 recognition motif to displace the regulatory sequence from the substrate binding cleft, enabling its capture by an adjacent subunit and R_1_ T287 trans-phosphorylation (41). Longer Linker segments extend the reach of “activation competent” subunits in auto-inhibited holoenzymes (42) to enable cooperative kinase activity by facilitating capture (41). The extended reach may also allow single holoenzymes to cross-link adjacent actin filaments or bind multiple actin subunits within the same filament (15). The 90 residue β_Rat_ linker, named “actin binding domain” (9), has an established role in actin binding; although isoforms with shorter linkers also bind actin (α_Rat_ ~25 residues (15)) and form F-actin networks (γ_Rat_ ~ 40 residues (38)). Multiple serine and threonine phosphorylation sites along the length of the β_Rat_ linker modulate CaMKII F-actin association (7), but they are not conserved across species. We assessed the role of linker length on network formation; using the structurally-characterized *C. elegans* CaMKII, “d” isoform (d_C.elegans_) (Chao et al., 2010) which has a 3-fold shorter linker than β_Rat_ which in turn has a 3-fold shorter linker than the human CaMKIIβ isoform (β_Hum_) (Bhattacharyya et al., 2016). Bioinformatic analysis predicted secondary structure for clues on the compliance of the linker and adjacent regions.

Finally, SPT measured the association of single GFP-β_Rat_ holoenzymes with synthetic F-actin assemblies and G-actin to relate their association / dissociation kinetics to network assembly / disassembly. Our measurements reveal that CaMKII cross-linked actin gels have sufficient mechanical resilience to maintain their basal network structure; but disassemble rapidly in response to a calcium-calmodulin pulse and are then remodelled by myosin motor forces with kinetics consistent with reported changes in stimulated spine volume.

## Materials & Methods

### Materials

Actin antibody specific to the C-terminus of α-actin, #A5060 (rabbit), was purchased from Sigma. Monoclonal GFP antibody, #1814460 (mouse) was purchased from Roche and the GFP protein, #8365-1, was from Clontech. A twenty-one-residue peptide, tatCN21, homologous to the NMDA type GluN2B receptor subunit (43), was a gift from Dr Ulli Bayer. Polyethylene glycol (MW 3KD (PEG)) was from Sigma. Buffer stocks for fluorescence assays were, AB^-^: 25 mM imidazole-HCl, 25 mM KCl, 1 mM EGTA, 4 mM MgCl_2_, pH 7.4. AB^+^: as AB^-^ but with additional 2 mM ATP. Ca.AB^-^ and Ca.AB^+^ contained 0.2 mM CaCl_2_ in place of 1 mM EGTA. Calmodulin, purified as detailed below, was used at 10 µM final concentration unless noted otherwise. Immediately prior to the experiments an oxygen scavenger system was added to 1-ml aliquots of the relevant, degassed, buffers to reduce photo-bleaching. This comprised: 20 mM DTT, 0.2 mg/ml of glucose oxidase, 0.5 mg/ml of catalase, 3 mg/ml of glucose, 0.5 mg/ml of bovine serum albumin (BSA) (final concentrations). PEG solutions were made in AB^-^ buffer with GOC. Protein expression and purification are detailed in **Supporting Material Section A**.

### Epi-fluorescence and total internal reflection fluoresce (TIRF) light microscopy

Epi-fluorescence microscopy assays studied rhodamine-phalloidin stabilized F-actin (“Rh-Ph F-actin”) gels and their motile response to crowding and motor forces. The assays used a Zeiss Axioskop 40, upright fluorescence microscope (44), equipped with: 100W mercury arc lamp; rhodamine filter set (Ex:525AF45, dichroic:560DRLP, Em:595AF60, Omega Optical, Brattleboro, VT, USA); 100x PlanNeofluor 1.3NA objective; PTI IC300 image-intensified CCD camera (Horiba, Kyoto, Japan). Camera magnification = 78 nm / pixel or nm/pixel (selected using a 1.5x magnification lens). Video records were captured using a frame grabber card (Picolo, Euresys, Multipix Imaging Ltd, Petersfield, Hampshire, UK) and recorded onto computer hard disc. Defined depth chambers were used to estimate GFP- β_rat_ concentrations in extracts, with known GFP solutions as calibration standard. Dual color, single molecule TIRF imaging studies were conducted using a Nikon Eclipse, TE 2000U inverted microscope described previously (15) with modifications. Sample preparation and TIRF modifications are detailed in **Supporting Material Section B**.

### Image Analysis

**Single Particles:** Single particle tracking (SPT) was performed with GMimPro and further analysed with Motility (45) software. Kymograph and colocalization analysis used ImageJ plugins. Details are in **Supporting Material Section B**.

**Networks**: The SOAX program based on SOACs (Stretching Open Active Contours) (46) was used for automated representation of the LM imaged networks (sampling distance, c = 8 pixels (0.625 µm)). SOAX gave a map of the coordinates of filament crossover points (“junctions”) and outlines of the connecting cables (“snakes”) superimposed on the original image **(Supporting Material Movie S1)**. Snakes included both single filaments and bundles that could not be distinguished in the diffraction-limited images. SOAX parameters were iteratively optimized for maximal superposition of the map with the imaged network. Manual editing eliminated spurious snakes and fused neighbouring junctions < 1.2c apart. Junction centroids and all inter-junction distances, *l_x_*, were determined from the map. Nearest neighbour distances, (*r_NN_*) were then obtained from the table of *l_x_* values using a custom script in R 3.3 (*https://www.r-project.org/*). The average near-neighbour distance, *R_NN_*, was computed from the observed number, N, of r_NN_ values, expressed in µm.

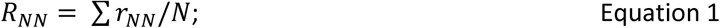

The mean snake intensity, *Q*, was determined in detector counts per image pixel area and mean snake curvature, K, measured in radians/µm was computed in SOAX from the tangent differences, ΔT, at user defined sampling distances, Δs, along the snake.

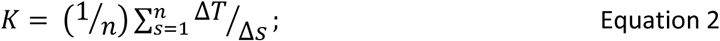

Calculation of the persistence length (*l_p_*) from *K* assumed a worm-like chain.

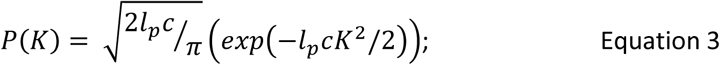

where *P*(*K*) is the probability density distribution for *K*, further details are in (47)

The *K* distributions were fitted well by the worm-like chain model however, *l_p_*, was systematically under-estimated (**Supporting Material Figure S1C**). Therefore, our quantitative analysis of network type was based mainly on *Q* and *R_NN_* with *l_p_* only as a qualitative check (**Figure 2A)**.

**Figure 2:**
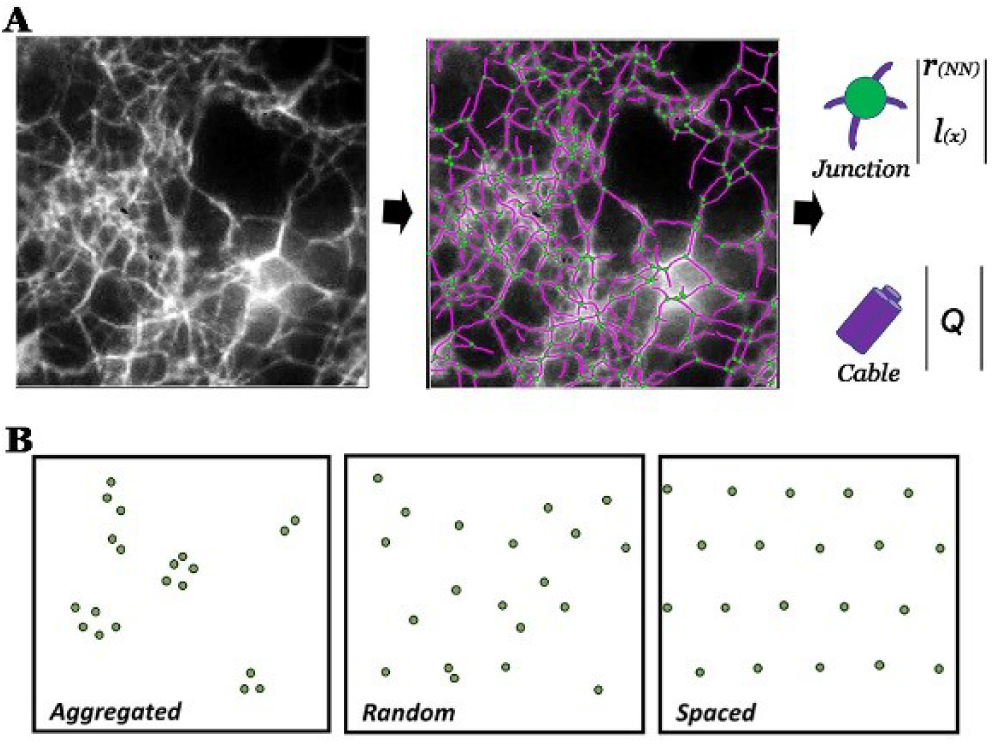
**A**. SOAX representation of a F-actin network in 15% PEG showing snakes (purple filaments) and junctions (green spots) superimposed on the image. The parameters extracted from the SOAX were the junction density, inter-junction separation l(x) and cable intensity (Q). The nearest neighbour distance (r_NN_) was computed from junction density and separation. **B**. Network architectures. Random network flanked by examples of aggregated and spaced networks.

For networks compacted by myosin motors, the fluorescence “foci” comprising conglomerated F-actin were identified from binarized images by setting area (>50 µm^2^) and intensity thresholds using the *Particle Analysis* function in ImageJ v1.5 (*https://imagej.nih.gov*). Foci centroid positions, inter-foci distances (*l_x_*) and nearest neighbour distances (*r_NN_*) were computed, as above for the snake junctions to obtain the integrated intensity, ∑*Q_foci_*.

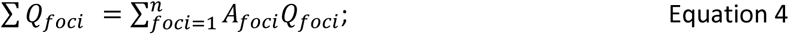

where *A_foci_* (pixel^2^) and *Q_foci_* counts / pixel are the area and mean intensity for each of *n* foci in an image field of view, using the binarized image to mask all foci located on the original image.

*R_NN_* was used to characterize the network type by comparison against the expected value for a random (Poisson) distribution, *r_rand_*, at a given density of junctions, *σ*, per unit area. It has been shown (48), that for a random distribution, obtained when the position of any point is equiprobable and independent of all other points:

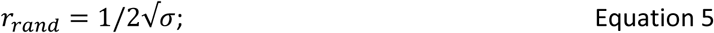

The limiting extreme divergences from the random distribution are: An aggregated distribution, where points coalesce and *R_NN_* approaches zero and an optimally spaced distribution on a hexagonal point spacing (**Figure 2B**) where 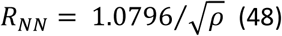 (48). Here, we define a “Randomness Index”, *R* = 221 *R_NN_*/*r_rand_* for our network architectures. *R* will vary from ~0 for the aggregated case to unity for a random (non-interacting) network up to 2.16 for the evenly-spaced hexagonal lattice.

## Results

### A. Assembly / disassembly of CaMKII-actin networks

#### 1. Imaging of CaMKII networks

CaMKII-actin assemblies were studied by negative stain electron microscopy (EM) and fluorescence light microscopy (LM). F-actin formed connected networks with single filaments and filament bundles connected by cross-over junctions in the presence of sub-stoichiometric β_Rat_ / G-actin molar ratios (0.3) (**Figure 3A**). The β_Rat_ holoenzymes localized preferentially at the junctions and along bundles. Examples of single holoenzymes at junctions connecting multiple filaments were common. Smaller AD hubs resulting from proteolysis did not bind. Both d_C.elegans_ and β_hum_ formed networks with similar porosity at comparable concentrations in spite of the large variation in linker length (25 (d_C.elegans_) −200 (β_hum_) residues). The increased β_hum_ linker length did not increase the number of holoenzymes per junction that rarely exceeded two in the EM images (**Supporting Material Section B Figure S2**). The architecture of networks formed by the proteins with Rh-Ph F-actin, imaged in solution with LM, was similar. The preponderance of cables relative to junctions increased dramatically as seen in the example of a network formed with d_C.elegans_ (**Figure 3Bi, Supporting Material Movie S2**) when concentration was increased three-fold. In conclusion, the nematode holoenzyme with short linkers forms networks almost as well as the long linker vertebrate holoenzymes at physiological molar ratios.

**Figure 3:**
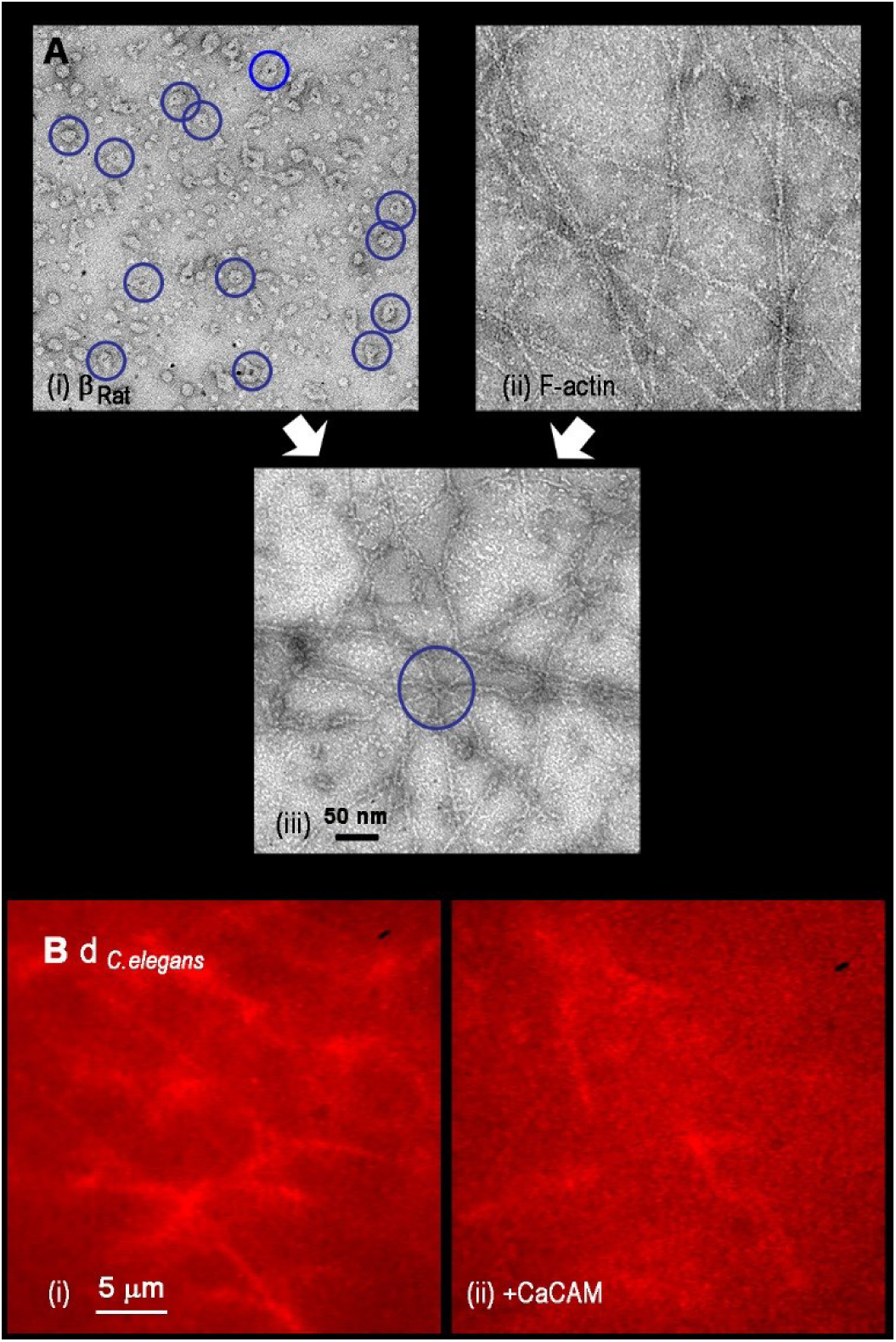
Architecture of CaMKII F-actin networks. **A. (i)** β_Rat_ holoenzymes (blue circles) and **(ii)** F-actin form **(iii)** a connected network. A single holoenzymes junction with multiple F-actin filaments (centre (blue circle)). **B (i)** Rh-Ph biotinylated F-actin gel with d_C.elegans_. on streptavidin coated glass after 30-minute incubation (Q =13030 counts / pixel, R = 0.75) and **(ii)** Disassembled filaments < 3 minutes after subsequent flow-in of 0.1 mM calcium / 1 µM calmodulin.

In all cases, the network disassembled when perfused with calcium-calmodulin (< 1 minute) as assessed by increased thermal motions and decreased intensity of the filaments (**Figure 3Bii, Supporting Material Movies S3**). The disassembly time is consistent with the loss of β_Rat_ co-localization with the neuronal F-actin cytoskeleton upon glutamate stimulation (18). It provided a diagnostic of CaMKII mediated F-actin assembly. In conclusion, the EM established that single CaMKII holoenzymes from three evolutionarily distant species whose linkers varied in length over a 9-fold range bound multiple actin filaments at junctions to form connected networks. The LM established that networks formed by the invertebrate d_C.elegans_ were disassembled rapidly by calcium-calmodulin.

#### 2. Bioinformatics of the actin binding domain

We analysed the linker sequences to understand the surprising independence of network assembly on linker length (**Supporting Material Section C**). Sequence homology between CaMKII isoforms / splice variants of the three species was obtained for two small linker peptides: a previously identified core fragment (β_Rat 342_KSLNNKKAD_351_ (39)) and another serine / threonine rich 20 residue peptide at the β_Rat_ linker / AD interface. Phylogenetic tree construction from the Uniprot CaMKII database revealed that the population diversity in both the KD and linker domains was sampled completely by the human CaMKII isoforms relative to the *C. elegans* splice variants that were confined to a branch within the trees (“monophyletic”). The increased diversity in the vertebrate populations may be a consequence of gene duplication. Linker length variation was a dominant factor in the greater spread of the linker versus the KD phylogenetic tree.

We used DisoPred (49) to identify unstructured (disordered) regions that may allow kinase domains to extend out from the hub. We found that the 90-residue linker together with about 40 N-terminal AD residues is the most unstructured part of the β_Rat_ protein. The shorter linker rat isoforms had similar disorder profiles over over-lapping sequence segments. Remarkably, the entire β_Hum_ linker segment subunit (>100 residues) additional to the β_Rat_ linker, was predicted to be unstructured. The known crystal structure (KD+AD) of the β_Hum_ subunit (with unknown linker structure) was used to check our secondary structure predictions and the structured regions were superimposed on the disorder profiles. An α-helix was predicted with high confidence for the core peptide, but the other conserved peptide was unstructured. The two peptides may bind actin directly, but the multiple phosphorylation of the unstructured peptide, as well as adjacent segments, must have a regulatory role (7). The conserved disorder profiles across isoforms and species reinforce the idea that linker compliance is an important determinant of its functions.

### B. Responses of CaMKII actin networks to mechanical stress

Calcium-calmodulin disassembled the CaMKII-actin networks but translational diffusion of the filaments was too slow to remodel the network over the sub-minute time scale relevant for early LTP. Therefore, we studied the role of additional forces that might operate within living cells. Specifically, we examined the resilience of CaMKII-actin networks, in the presence of myosin motor activity and molecular crowding both in the absence and presence of calcium-calmodulin.

#### 1. Macromolecular crowding by globular solutes

We first examined crowding by polyethylene glycol PEG, an inert globular solute, whose effects on F-actin organization have been studied previously (33) (**Figure 4A**). The SOAX snake intensity (Q) distribution mapped the increase in bundling with PEG concentration (**Figure 4B)**. The persistence length (*l_p_*) computed from snake curvature (*K*) increased (i.e. the filaments stiffened) as the networks formed; from 6 µm (10% PEG) to 10 µm (15% PEG) with little significant increase at higher PEG concentrations. Bundling increased with PEG concentration; with a sharp increase between 15-20% as aggregated filaments aligned into bundles. The estimated viscosity of 20% PEG solution (8.6 cp) (**Supporting Material Figure S1B**) is comparable to estimates for dendritic spine cytoplasm (50). The transition to the aggregated state took over 20 minutes to complete, as tracked by Q and R with F-actin alone (**Figure 4C**) or with β_Hum_ in calcium-calmodulin 20% PEG buffers. CaMKII-actin networks formed with β_Hum_ or β_Rat_ were insensitive to crowding by PEG in absence of calcium-calmodulin (**Figure 4D**). We noted that the increased linker length of the β_Hum_ isoform failed to promote formation of multiple-filament junctions and stronger networks as one might perhaps have expected if the longer linker were to allow more flexible geometry of binding and increased F-actin cross-linking.

**Figure 4:**
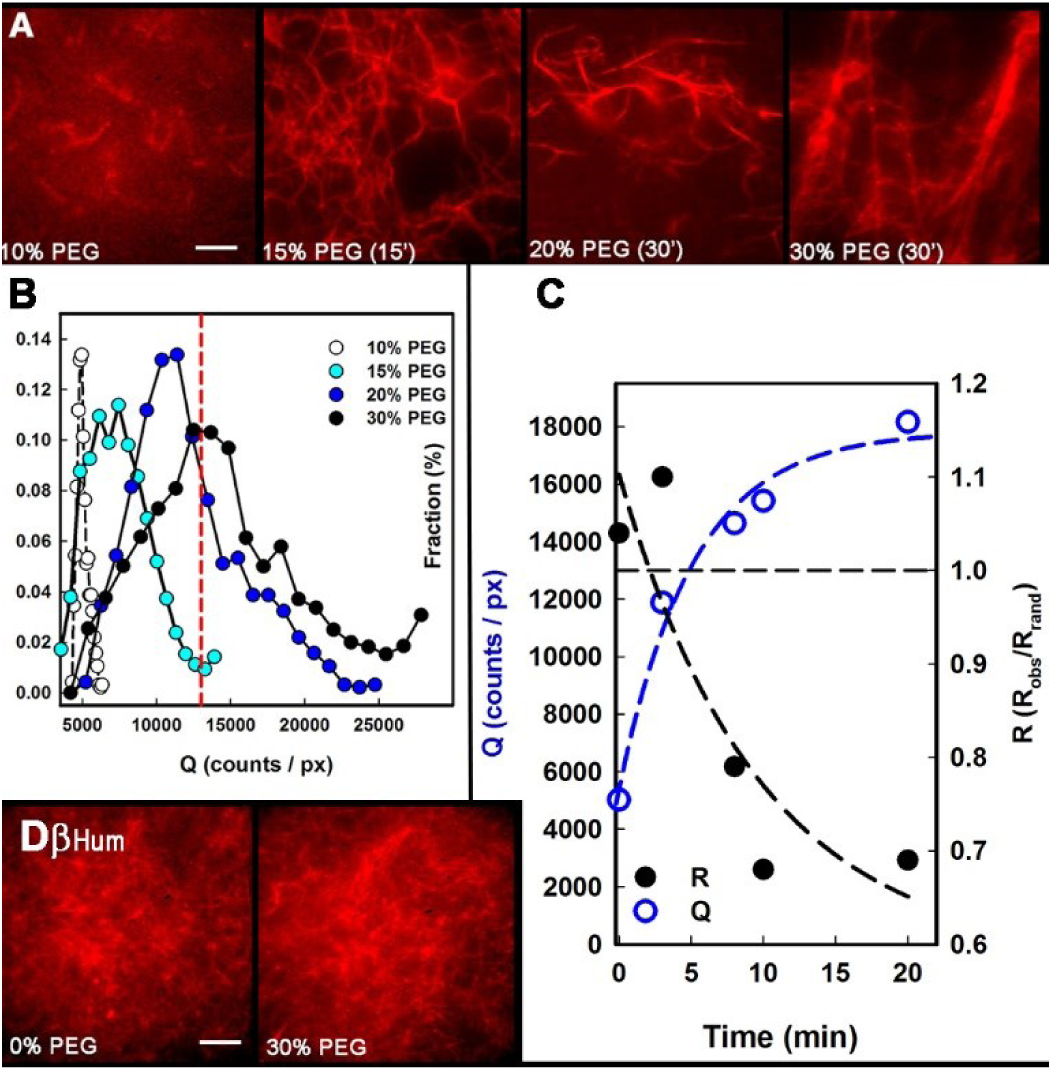
CaMKII counteracts macromolecular crowding. **A**. Rh-Ph F-actin suspensions in AB^-^/GOC with 0% PEG, 15% PEG (15 minutes), 20% PEG (30 minutes) and 30% PEG (30 minutes). **B**. Cable intensity (Q) distributions as a function of PEG concentration show shifts in modal values as bundling increases. Dashed line (red) is Q value for d_C.elegans_ network (Figure 3Bi). **C**. Time course of network formation (20% PEG calcium-calmodulin buffer, F-actin /β_Hum_) tracked by the increase in Q, decrease in R as the 20% PEG network forms. Dashed lines show single exponential best-fits. **D**. Rh-Ph F-actin/β_Hum_ mixtures in 0% and 30% PEG (R = 1.04, Q = 7810 counts / pixel). Scale bar (A & D) = 5 µm.

#### 2. Network remodelling by myosin motors

In order to study the ability of myosin-II motors to reorganize and/or compact CaMKII-actin filament networks we employed two different geometries: In the first, we coated the microscope coverslip surface with HMM to form an isotropic layer of myosin motors, using procedures adopted for in vitro motility assays. In the second format, we attached CaMKII-actin networks to the coverslip surface using biotin-streptavidin linkages and then introduced myosin mini-filaments into the bulk volume of the networks.

##### (a) Surface-immobilised myosin motors

In the absence of ATP, mixtures of d_C.elegans_ CaMKII and F-actin (at 1:3 molar ratios) formed a contiguous, interconnected filament network that attached to HMM-coated surfaces. The network did not disassemble when ATP (AB^+^) was perfused in (**Figure 5 A, B**). A sub-population of filaments that was not part of the network aligned (51) and glided along rails of static filaments in the immobilized network (**Supporting Material Movie S4**). Fragmentation upon ATP addition, a common occurrence at high HMM densities (52), did not occur as the static filaments buttressed against orthogonal motor forces on the motile filaments to promote gliding over fragmentation. Furthermore, the CaMKII-actin network did not alter upon perfusion of concentrated (1.2 mg / ml) mixtures of non-fluorescent, phalloidin-stabilized F-actin. However, when calcium-calmodulin was introduced networks disassembled over a period of ten minutes and dissipated as the HMM translocated the separated filaments. F-actin networks were not created if the CaMKII / F-actin mixture was prepared in the presence of calcium-calmodulin. In this case, filaments glided upon addition of ATP and formed patterns upon perfusion of the unlabelled F-actin solutions as seen before (44).

**Figure 5:**
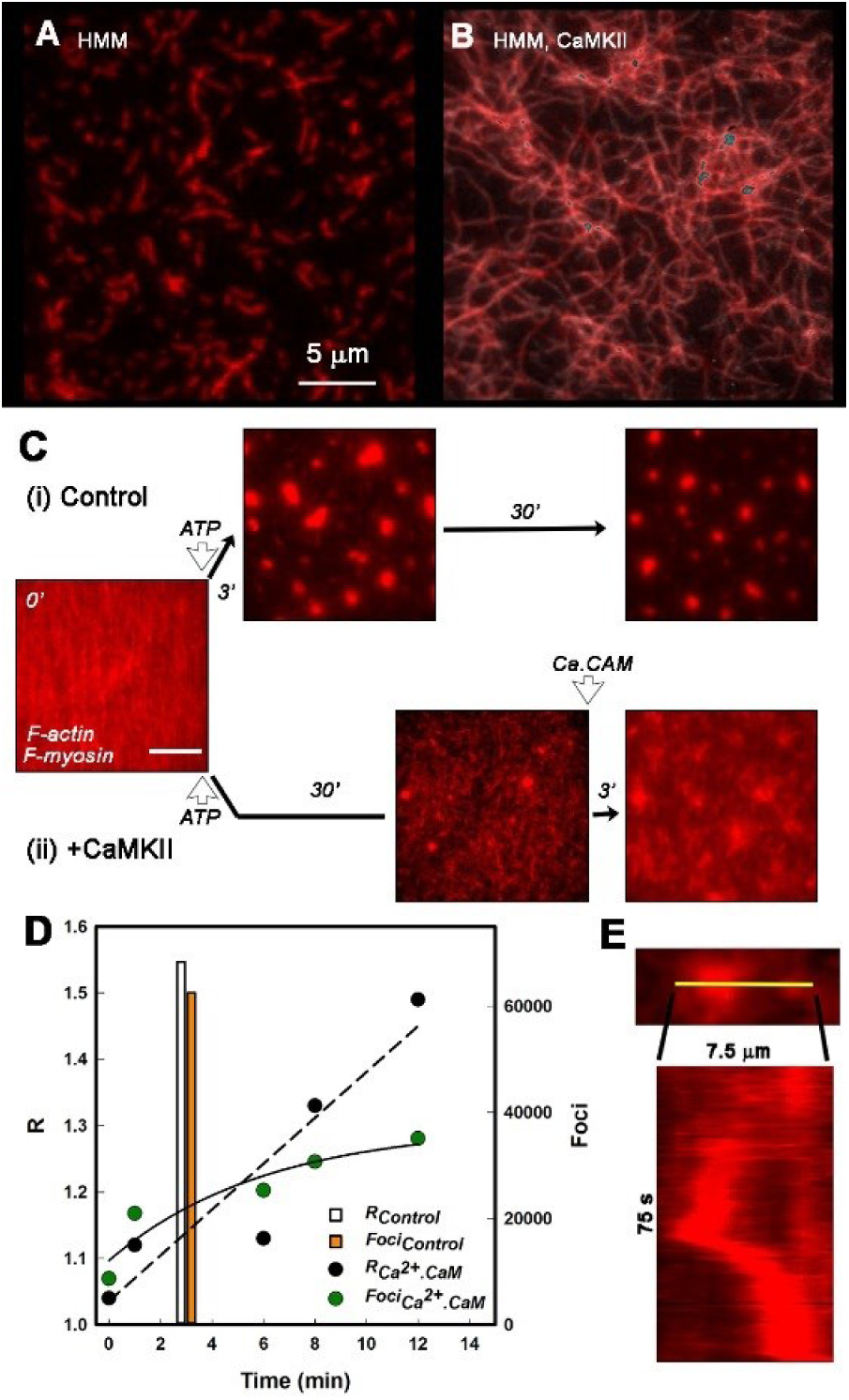
Long-range alignment by myosin motor coated surfaces and myosin filaments. **A**. F-actin filaments ((l_x_) = 1.5 µm) gliding on a HMM coated surface. The gliding filaments align when crowded. **B**. F-actin filaments gliding (silver outlines) on a HMM coated surface along immobilized filaments (red). ((l_x_)= 2.5 µm) cross-linked by d_C.elegans_. Gliding was initiated after 30-minute incubation on HMM coated glass by flow-in of AB^+^/GOC. Scale as for A. **C**. Rh-Ph biotinylated F-actin tethered to streptavidin coated glass coverslips incubated with myosin filaments. Compaction after perfusion of AB^+^/GOC (2 mM ATP) in the absence **(i**) and presence of **(ii)** d_C.elegans_. After perfusion of calcium buffer Ca-AB^+^/GOC with 1 µM calmodulin the CaMKII cross-linked network changes to resemble the control network. All scale bars (white) = 5 µm. **D**. Time course of network remodelling after calcium-calmodulin perfusion tracked as an increase in R (black) and ∑Q_foci_ ((Foci) green) compared to the time for foci formation (R (white), ∑Q_foci_ (orange) in the absence of CaMKII. Linear (dashed line) and exponential (continuous curve) best-fits to R and Q respectively are shown. **E**. Kymograph of the approach and coalescence trajectories of two foci (from a central section of the image field in Movie S6) along the yellow line.

##### (b) Bulk myosin mini-filaments

Next, we tested the effect of concerted motor forces exerted by myosin mini-filaments on the mechanical resilience of CaMKII-actin networks. Surface-tethered, 3D CaMKII-actin networks were created by first adding a layer of biotinylated F-actin (1:10 biotin / G-actin labelling ratio) to a neutravidin-coated coverslip followed by perfusion of different mixtures of non-biotinylated Rh-Ph F-actin, myosin mini-filaments and d_C.elegans_ CaMKII in AB^-^. After 30-minute incubation any unbound material was washed away with AB^-^ and networks viewed by epi-fluorescence microscopy. First, a control experiment, performed in absence of CaMKII, showed that ATP addition (AB^+^) triggered rapid compaction of the actin filaments (**Figure 5C, Supporting Material Movie S5**) as reported (36). Fluorescent F-actin foci coalesced (< 3 minutes) and compacted until the force balance between myosin filaments at different foci caused them to become spaced and pulled the inter-connecting F-actin cables taut. Next, F-actin networks were formed in the presence of CaMKII. Now, compaction was largely blocked over a 30-minute period after addition of ATP. However, subsequent addition of calcium-calmodulin initiated rapid compaction that was >50% complete within 3 minutes **(Supporting Material Movie S6)**. After six minutes additional foci did not form but intensity of existing foci and the spacing between them increased. The “R” value increased linearly over a 12-minute period, (i.e. the foci spacing became more ordered with time) while the integrated foci intensity, ∑*Q_foci_* showed a single exponential increase to a limiting value (**Figure 5D**). Coalescence dynamics of two adjacent foci with brief episodes of rapid translocation over a 75 second period (0.1 µm / s approach rate) are shown in an example kymograph (**Figure 5E**).

### C. Association / dissociation of single holoenzymes with F-actin and G-actin.

We used single molecule TIRFM to test for differential attachment of the GFP-β_rat_ holoenzymes with network junctions and cables predicted from the EM and LM analysis of the networks. We also characterized their dissociation by calcium-calmodulin.

#### 1. F-actin

When viewed using TIRFM individual GFP-β_rat_ holoenzymes (10 - 50 nM concentration) appeared as diffraction limited spots which decorated immobilized synthetic Rh-Ph F-actin filaments (**Figure 6A-C, Supporting Material Movie S7**). The spots localized along single filaments and filament bundles (distinguished by snake intensity Q value), and preferentially at filament intersections (“junctions”). When a mixture of calcium-calmodulin and GFP-β_rat_ (in AB^-^ buffer) was perfused into the flow-cell, filament decoration was substantially reduced within ~20 seconds, the time taken to complete the buffer exchange. Thus, calcium-calmodulin rapidly dissociated the GFP-β_rat_ in the absence of ATP. A residual GFP-β_rat_ fraction remained bound to F-actin and this fraction was unaffected by subsequent addition of ATP (calcium-calmodulin AB^+^ buffer). These results were consistent with the observation that calcium-calmodulin disassembled CaMKII-actin networks in the absence of ATP (Figure 3B, above).

**Figure 6:**
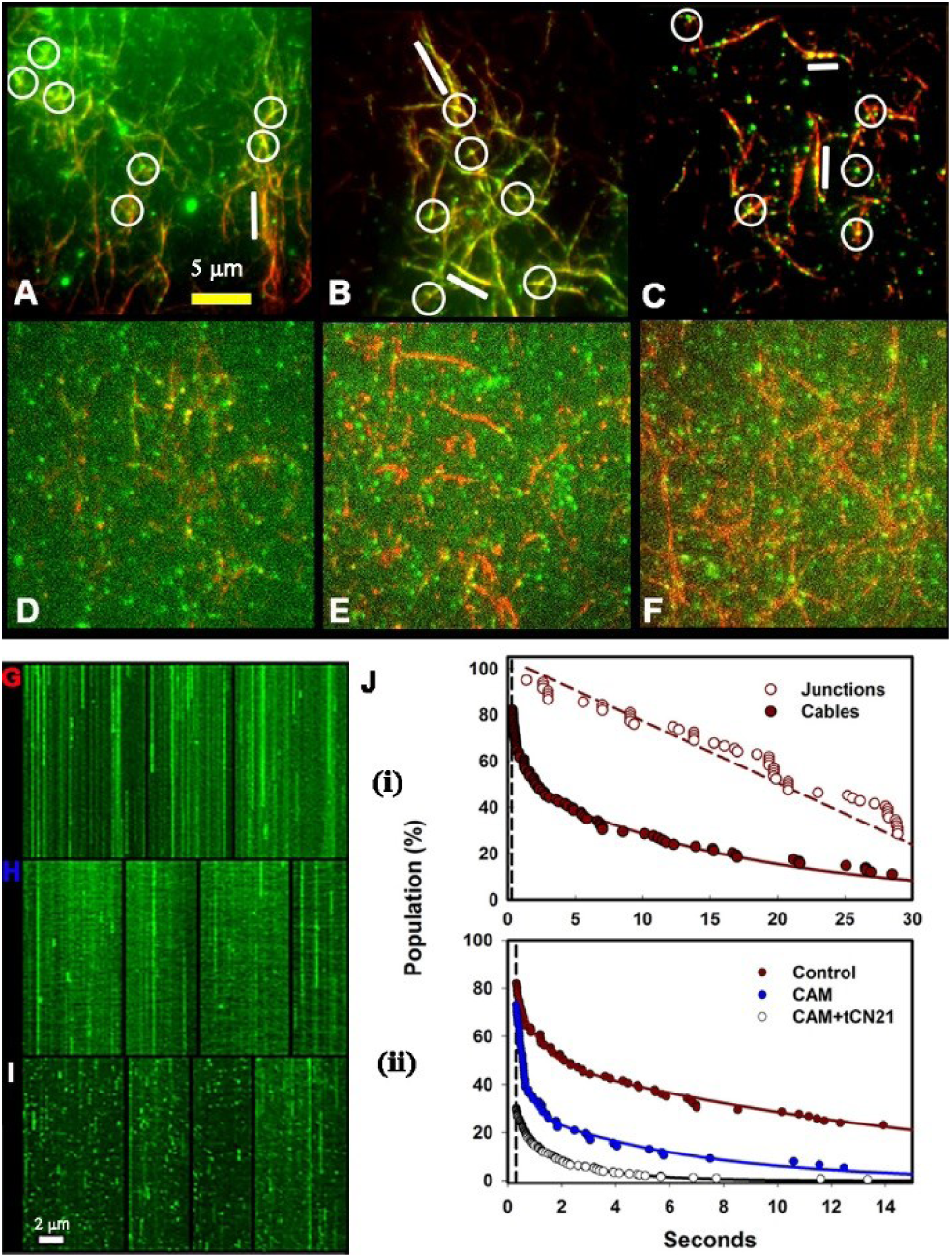
Single molecule binding to F-actin filaments. Averaged dual colour TIRF of GFP-β_Rat_ holoenzymes (green spots) bound to single Rh-Ph F-actin (red filaments). 20 frames / second video frame rates. **A, B, C**. AB^-^/GOC buffer. frame averages. Circles (white) mark GFP-β_Rat_; bars (white) mark filament stretches decorated continuously with GFP-β_Rat_. **D, E, F**, AB^-^/GOC buffer with 1 µM calmodulin / 0.2 mM calcium. 2 frame averages. Scale bar for panels A-F (yellow). **G, H, I**. Kymographs of green fluorescent spots associated with actin filaments in **(G)** absence and **(H)** presence of calcium-calmodulin or **(I)** calcium-calmodulin plus tCN21. The duration (abscissa) of each kymograph is 20 seconds (400 frames). Bound fractions (dwell time > 6 frames (0.3 s)) are 0.82±0.11 (**G**), 0.73±0.09 (**H**), 0.30±0.01 (**I**). Length scale (white) = 2 µm. **J**. Dwell-time distributions of the three conditions, colour coded to match panel labels. k_off_’s were determined from single or double-exponential best-fits (solid lines). Vertical dashed lines mark 6-frame “bound” spot threshold **(i)** Comparison of the distributions of control sub-populations localized at junctions (empty red; k_off_ = 0.04 s^-1^) or cables (solid red; rate = 0.43(exp (−1.2) + 0.56(exp (−0.06) s^-1^). **(ii)** Distributions of bound control (red), calcium-calmodulin (blue; rate = 0.78(exp (−3.1) + 0.22(exp (−1.4) s^-1^) and calcium-calmodulin (+ tCN21) (white; k_off_ = 1.0 s^-1^) treated populations.

We then used tCN21, as an analogue of the NMDA receptor, to explore whether the CaMKII-Receptor interaction would affect the residual CaMKII-actin association. We found that the residual bound fraction (i.e. the fraction of F-actin bound CaMKII molecules resistant to calcium-calmodulin) was reduced when tCN21 was perfused with GFP-β_rat_ in calcium-calmodulin AB^-^. We also found an increase in CaMKII-actin dissociation rate (reduced dwell-time) (**Figure 6D-F, Supporting Material Movie S8**). In fact, measurable binding was only detected when three-fold higher GFP-β_rat_ concentrations relative to those used earlier were perfused in with tCN21. We speculate that the residual GFP-β_rat_ fraction represents the compact autoinhibited form inaccessible to calcium-calmodulin (53). tCN21 binding to conformations opened by calcium calmodulin shifts the steady-state distribution between the compact and extended auto-inhibited forms towards the latter, allowing more calcium-calmodulin to bind and dissociate GFP-β_rat_ holoenzymes from the F-actin.

Kymographs (**Figure 6H-I**) analysed dwell times of GFP-β_rat_ holoenzymes. The particles rapidly exit the evanescent field once dissociated from binding targets as detailed in an earlier publication (15). The analysis (**Figure 6J**) revealed that junction-localized CaMKII had substantially longer dwell times (slower “off-rate (*k_off_*)”) than the rest of the population implying that the steady-state observation of preferential CaMKII localization at junctions was due to slower dissociation. The rotationally symmetric CaMKII holoenzyme can readily attach to multiple filaments congregated at cross-over junctions. Dissociation by detachment from all filaments at a junction should take longer than termination of single attachments that will mostly form with isolated filaments. The mean intensity of the junction sub-population was not measurably greater than the mean of the population localized at other points along F-actin consistent with the EM evidence that junctions are predominantly formed by single holoenzymes.

#### 2. G-actin

We tested the association of Cy3 conjugated G-actin (Cy3-actin) and GFP- β_rat_ for estimation of monovalent dwell times. Glass coverslips coated with antibody against G-actin or GFP were prepared. In one geometry Cy3-actin was immobilized on actin antibody-coated glass. GFP- β_rat_ formed a dense layer associated with the immobilized G-actin spots when perfused in (**Figure 7A**). The SPT lifetime distribution was well-fit by a single exponential with rate, *k_off_* = 2.55 s^-1^ (n > 1200). In the presence of calcium-calmodulin, there was a dramatic decrease in the number of GFP- β_rat_ spots in the evanescent field adjacent to the immobilized Cy3-actin surface layer with high background fluorescence due to distant particles outside the field. However, a few GFP- β_rat_ particles rapidly diffused in and out of the evanescent field with brief stationary episodes in proximity to G-actin spots as realized by kymograph analysis (n = 12). An example kymograph is shown (**Figure 7B**). The dramatic difference in the evanescent fluorescence and its spatiotemporal dynamics between the two cases is best appreciated from the video-records (**Supporting Material Movies S9, S10**).

**Figure 7:**
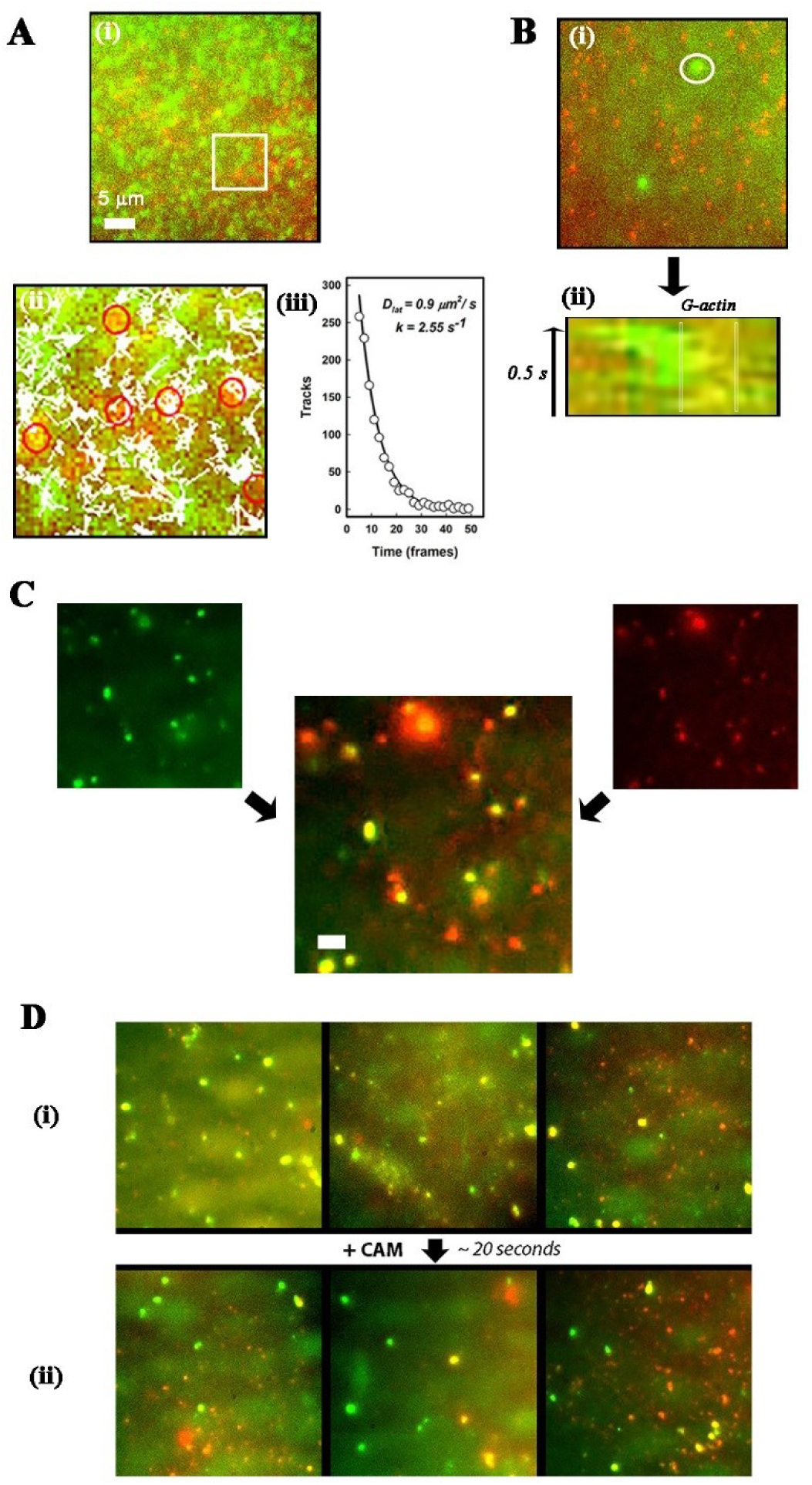
**A (i)** GFP-βrat (green) molecules in the evanescent field adjacent to a surface layer of antibody immobilized Cy3 G-actin (red) molecules. **(ii)** Particle tracks superimposed on snapshot of part of the image field (Box area in (i)). **(iii)** Distribution of track lifetimes fits a single exponential decay (k = 2.55s^-1^, R>0.99). **B (i)** Experimental conditions as in (i), but in presence of calcium-calmodulin. **(ii)** Kymograph (circle in (ii) 3 µm diameter) shows transient interaction of GFP-βrat particle with a Cy3-actin molecule (demarcated). **C**. Colocalization of Cy3 G-actin with antibody immobilized βrat molecules. 50-frame average. Scale Bar (white) = 2 µm. **D**. Image fields from an experiment comparing Cy3 G-actin co-localization in the absence (**(i)** 0.33 ± 0.08 Pearson’s correlation)) and presence (**(ii)** 0.015 ± 0.04 Pearson’s correlation)) of calmodulin. The latter images (50 frame average) were recorded immediately after introduction of calmodulin.

Alternatively, GFP antibody-coated glass coverslips trapped GFP- β_rat_ molecules from the extracts. The GFP- β_rat_ mostly attached as diffraction limited spots that remained when the extract was washed out. When solutions containing Cy3-actin were perfused through the flow cell the Cy3-actin co-localized with a dominant fraction of the GFP spots (**Figure 7C**). Perfusion of calmodulin in the calcium AB^-^ buffer triggered a large drop in the occupied fraction that was complete (< 20 seconds) before acquisition of the video-record (**Figure 7D**). A minor fraction (~ 20%) of the co-localized Cy3-actin was refractory to dissociation by calmodulin, reminiscent of the residual binding of GFP- β_rat_ to F-actin.

## Discussion

In the present study, four aspects of CaMKII-actin networks have been explored; **(1)** Assembly and mechanics of CaMKII cross-linked F-actin networks; **(2)** Physical role of the CaMKII linker region in network connectivity; **(3)** Modulation of actin affinity by calcium-calmodulin, ATP and tatCN21; **(4)** The impact of molecular crowding agents and myosin motors on network architecture. Our results contribute to our understanding of the structural role of CaMKII in dendritic spines. The implications of our results for the structural role of CaMKII in dendritic spines and spine remodelling during early LTP (**Figure 1**) are discussed.

### 1. Assembly and mechanics of cross-linked actin networks.

Network architecture was characterized based on two parameters; cable thickness (reported by the value, Q, which for the diffraction-limited case increases with bundle size) and nearest-neighbour junction distance R_NN_ compared to that expected for a random network, R_rand_. The ratio (R_NN_/R_rand_) was used to compute a randomness parameter “R” reporting the orderliness of the filament junctions. Observed changes of these parameters described F-actin network architectures under both steady-state and dynamic experimental conditions. A random network with Poisson distributed nearest neighbour junction distances (R~1) was obtained at low CaMKII-actin molar ratios implying little or no interaction between neighbouring nodes. The network was stabilized by preferential localization of CaMKII to junctions. When the ratio of CaMKII to F-actin was increased, F-actin bundles were progressively formed and network architecture transitioned to a non-random, aggregated, type (R<1). The molecular crowding agent, PEG, caused a similar transition by causing F-actin bundling and collapse of the random network.

EM images of β_Rat_.F-actin show that junctions are abundant at sub-stoichiometric (0.3) ratios. Published electron micrographs of CaMKII-actin networks show images similar in appearance to ours at comparable molar ratio; while bundling dominates (> 80%) at equimolar and higher ratios (9). The LM and EM make the case that filaments are stapled together by the holoenzymes in stochastic, polarity-insensitive fashion and explain the absence of any periodicity associated with CaMKII bundles in studies thus far, in contrast to other actin binding proteins (54).

A study (36) of streptavidin / F-actin_biotin_ cross-linked networks mixed with myosin filaments showed that upon ATP addition and myosin filament force production, streptavidin cross-links increased the coalescence of actomyosin foci (0.6 µm/s) from the rate in absence of streptavidin (0.1 µm/s) at low biotin levels (1 / 1000 biotin / actin molar ratio), but retarded (0.03 µm/s) at 5-fold higher levels due to tension of connecting filament cables opposing motor forces. Here, we found that coalescence of the foci was blocked in the presence of CaMKII. We conclude that under our experimental conditions, CaMKII-actin networks have sufficient mechanical resilience to resist the compaction forces exerted by myosin mini-filaments. We note that 3-fold more actin filaments can be connected at a junction formed by a single CaMKII dodecamer relative to one formed by a streptavidin molecule with biotinylated F-actin, although a biotin-streptavidin bond has greater strength. The rate of coalescence upon disassembly by calcium-calmodulin was comparable to the reported control rate (36), indicating that any residual CaMKII-actin attachments do not impede the mechanics.

CaMKII-actin networks can complement studies of model systems such as streptavidin-biotin cross-linked actin networks in important ways by exploitation of calcium-calmodulin as a tool to dial the affinity and number of cross-links formed. The randomness parameter introduced in this study provides a ground-level descriptor of model networks to frame theories regarding their assembly and mechanism. It would be of great interest to relate this measure to higher level descriptors of bundle mechanics and viscoelasticity (37, 55) in future work.

### 2. The role of the linker domain in CaMKII-actin assemblies

The networks documented for all three CaMKII species in this study extends previous evidence from rat isoforms (38) that much of the β_Rat_ actin binding linker domain is not essential for F-actin association. The long-linker β_Hum_ forms networks indistinguishable from those formed by β_Rat_ at similar concentrations. The nematode d_C.elegans_, whose linker has comparable length and sequence to the α_Rat_ and γ_Rat_ linkers, also forms networks albeit at higher concentrations. Rapid-freeze replica images show extensible segments in the α_Rat_ linker (56). The persistence length of unstructured peptides is short (~ 1 nm) (57) so even kinase subunits with short linkers may flex and bind multiple F-actin subunits. The d_C.elegans_ data document the F-actin association and its calcium-calmodulin triggered dissociation in invertebrates for the first time to our knowledge to underline their broad consequences for cytoskeletal signal transduction across evolutionarily distant species separate from the vertebrate cardiovascular and neuronal systems that have driven CaMKII research thus far.

The two conserved segments, the predicted α-helix core piece and an unstructured serine / threonine rich sequence element may directly bind actin; although the possibility that the actin binding residues are outside the linker (38) cannot be ruled out. Serine / threonine phosphorylation increase persistence length of unstructured peptides (58) and this may explain why mutation of all auto-phosphorylation sites to aspartate, a phospho-threonine mimic, in the linker and adjacent residues abolishes F-actin binding; but mutation to the small amino acid alanine has a small effect (7). The bioinformatics and the mutagenesis argue that the conserved unstructured peptide could be a regulated, flexible connector; consistent with growing evidence that unstructured peptides are prominent in signalling mediated by phosphorylation regulated protein-protein interactions (59).

The β_Hum_ linker does not increase actin network connectivity though remarkably, most (>90%) of its 220-residue sequence is predicted to be unstructured. Linker flexibility has an established role in the cooperativity of CaMKII activation by calcium calmodulin (53, 60), a metric of conformational spread. So, linker length might optimally tune CaMKII for its response to calcium stimuli in different cellular contexts and different organisms (61). Cooperativity during activation has been studied thus far with short (< 40 residues) linkers. In the future it would be of interest to investigate whether the β_Hum_ linker increases activation cooperativity.

### 3. Mechanism of calcium-calmodulin triggered dissociation.

The single molecule evidence for direct GFP-β_Rat_ binding to G-actin and F-actin extends conclusions from less direct methods based on intact cell cytoskeletons (15, 18) or correlated fluorescence intensity fluctuations (17). We found that calcium-calmodulin triggered GFP-β_Rat_ dissociation from G and F-actin that was too rapid to be measured and ATP was not required. The measured dwell-times showed associations – dissociations occur on the second time-scale for most (> 90%) of the population. The single-exponential fit for track lifetimes obtained for GFP-β_Rat_ association with G-actin contrasts with the tailed lifetime distributions, also obtained using live-cell TIRFM imaging and SPT, for GFP-β_Rat_ association at actin stress fibres in human umbilical endothelial cells (HUVECs) (15). We suggest that long-lived GFP-β_Rat_ holoenzymes trapped at filament junctions or bundles, as shown in this study, account for this difference. The long-lived, junction-localized fraction probably also limits disassembly (~ 1 minute) of the in-vitro β_Rat_ / F-actin networks.

A refractory sub-population that was not dissociated by calcium-calmodulin alone; was dissociated when the calcium-calmodulin was supplemented by tCN21. The peptide may have shifted the autoinhibited conformational ensemble towards extended linker states, visualized recently (62), in which the IQ-10 motif is accessible to calcium-calmodulin. In contrast, autocamtide-2 inhibitor peptide, an auto-phosphorylation site mimic, did not affect dissociation (7). The *k_off_* (inverse dwell-time) of the particle populations treated with calcium-calmodulin approached that measured by TIRFM ASPT for mutant T287D (6.5 s^-1^) GFP-β_rat_ that does not associate with cytoskeletal actin when expressed in the HUVEC cultures. All aspects listed above applied equally to CaMKII interactions with both G- and F-actin consistent with a common binding surface.

### 4. Implications for early LTP.

Following synaptic stimulation, cytosolic calcium increases rapidly to millimolar levels due to the inward NMDA calcium current to initiate the transition from **state (i)** to **state (ii) (Figure 1)**. Rapid (few seconds) dissociation of CaMKII from both F-actin and G-actin, as reported here, has a two-fold effect: Polymerization of the released G-actin that can proceed immediately and slower disassembly of the CaMKII F-actin network. The latter would coincide with the time course of myosin II polymerization into mini-filaments that would be slow as phosphorylation must precede myosin filament assembly. So, the rapid initial phase of spine expansion is due primarily to actin polymerization; while subsequent spine compaction occurs as actin polymerization slows and myosin motor activity increases. Although the calcium transient is short-lived the newly formed myosin mini-filaments as well as the PSD sequestered, NMDA bound CaMKII will continue to be active (63, 64).

The molar ratio of CaMKII / G-actin in the basal state of hippocampal neuron dendritic spines is around 0.3 (**Figure 1 – state (i))**. This estimate is based on the CaMKII concentration (138 µM) (10) and assumes actin is comparable to lamellipodia; µM as F-actin out of µM total (65). We find that <25% of F-actin is bundled at this molar ratio; but bundling occurs at higher molar ratios. The basal dendritic spine free-calcium level is sub-nanomolar so calmodulin is in its apo-form and CaMKII-actin networks are stable and not susceptible to forces due to macromolecular crowding or episodic myosin motor activation triggered by miniature synaptic currents (66). It is not presently known whether myosin motors act as individual molecules tethered to the PSD or as a collective by assembling into mini-filaments. Our assays recreated both scenarios with myosin motors at higher concentrations than likely to obtain at the synapse. Myosin II activity at the leading edge is thought to fragment actin networks for turning in neuronal growth cones (67). Neurons express the β_E’_ splice variant that lacks actin binding functionality during early development (16); so immature spine filopodia may also utilize myosin induced fragmentation unconstrained by CaMKII cross-links to orient to developmental cues. Fragmentation will, however, be deleterious for mature spines once synapses form so CaMKII-actin networks may be important blockers of episodic myosin activity.

The stable end state (**Figure 1 state (Iii)**) for spine volume is reached after 3 minutes. The kinetics of myosin powered remodelling, documented in this study, indicate that this is ample time for myosin action to be completed. On the other hand, compaction due to clustering mediated by crowding agents may be ruled out as it is too slow. As the new basal state is established the actin cytoskeleton will again be stabilized by CaMKII cross-linking. The tCN21 data suggest the CaMKII associated with NMDA receptors is mechanically uncoupled from cytoskeletal actin. The uncoupling may prolong PSD-sequestered CaMKII activation to allow PSD remodelling to be completed. However, CaMKII interactions with other cytoskeletal proteins, notably α-actinin (68), might be as important, if not more so, to synchronize the changes in cytoskeletal and PSD architecture for increased synaptic strength.

In conclusion, this study provides a novel view of CaMKII-actin dynamic architecture and evolution of the CaMKII actin binding modality. Measurements of the calcium-calmodulin triggered network disassembly and its modulation by ATP and membrane receptor targets have mechanistic implications that may be tested and refined by numerical simulations and in vivo-experiments.

## Abbreviations

EM: electron microscopy
LM: light microscopy
TIRF: total internal reflection fluorescence
CaMKII-actin: CaMKII / F-actin complexes
Rh-Ph F-actin: rhodamine phalloidin stabilized actin
calcium-calmodulin: calcium bound calmodulin

## AUTHOR CONTRIBUTIONS

S.K – Designed / performed experiments, analysed data, wrote manuscript. K.H.D – Designed / participated in EM experiments, edited manuscript. J.E.M – Designed experiments, analysed data, wrote manuscript.

## ACKNOWLEDGMENTS

We acknowledge Dr Margaret Stratton for gifts of bacterial CaMKII expression plasmids and advice on CaMKII purification protocols. This study was supported by grants from the Royal Collaborative Exchange (grant U1175), the Molecular Biology Consortium (S.K) and the Francis Crick Institute which receives its core funding from Cancer Research UK, the UK Medical Research Council, and the Wellcome Trust (J.E.M.). Electron microscopy was performed with equipment supported by NIH grant GM51487.

